# The proteomic landscape of genotoxic stress-induced micronuclei

**DOI:** 10.1101/2023.08.27.555027

**Authors:** Kate M. MacDonald, Shahbaz Khan, Thomas Kislinger, Shane M. Harding

## Abstract

Micronuclei (MN) are induced by various genotoxic stressors and amass nuclear- and cytoplasmic-resident proteins, priming the cell for MN-driven signalling cascades. Here, we measure the proteome of micronuclear, cytoplasmic, and nuclear fractions from human cells exposed to a panel of six genotoxins, comprehensively profiling their MN protein landscape. We find that MN assemble a proteome distinct from both surrounding cytoplasm and parental nuclei, with a core composition that is independent of the specific inciting stressor. Across stress conditions, MN are significantly depleted for spliceosome machinery and replication stress response proteins, but are enriched for a subset of the replisome. We find that the loss of splicing machinery within transcriptionally active MN contributes to intra-MN DNA damage, a known precursor to chromothripsis. This dataset represents a unique resource detailing the proteomic landscape of MN, guiding mechanistic studies of MN generation and MN-associated outcomes of genotoxic stress.

## INTRODUCTION

Micronuclei (MN) are cytoplasm-resident subcellular structures containing fragments of nuclear DNA, histones, and nuclear proteins, encased within a version of the nuclear envelope (1–3). MN arise infrequently in healthy cells, and are induced by unresolved DNA lesions or mitotic errors when cells are exposed to exogenous genotoxins (3,4). MN can actively drive a number of carcinogenic processes, including inflammatory signalling (3–6) and chromothriptic gene rearrangements (7–9) but these consequences are strongly context-dependent, and particularly sensitive to MN protein content. For example, MN initiate inflammatory cascades by recruiting the viral pattern recognition receptor cyclic GMP-AMP synthase (cGAS) from the cytoplasm (5,6). However, cGAS-MN interactions are maximized by integrity failure of the MN envelope, termed “rupture,” and rupture dynamics are determined by the protein composition of the micronuclear envelope (1–3,6). Upon rupture, cGAS-MN recruitment and activation are further influenced by characteristics of the enclosed DNA fragment (3,4,10) and by other proteins that may be present in MN, including exonucleases (11), transcription machinery (3), and translation cofactors (12). MN-driven chromothripsis is similarly influenced by MN content: Chromothriptic rearrangements are more likely when MN contain R-loops (7), replication machinery (8,13), and when they recruit certain DNA damage repair proteins once in the cytoplasm (7). Despite its importance, our understanding of the global micronuclear protein landscape is limited, and no datasets are publicly available for exploration. MN originate in the nucleus and reside in the cytoplasm, and it has not been demonstrated whether MN are assembling their own distinct, biased protein profile from each of these protein pools or, alternatively, passively acquiring high-abundance proteins from the nucleus at formation or from the cytoplasm once ruptured. Even among proteins generally accepted as MN-associated, such as cGAS, the strength of the association depends on the specific genotoxic stress condition used to generate MN (3,4). Beyond individual proteins like cGAS, the scale of the influence that a particular stress context has on the global MN protein landscape has not been defined. Addressing these open questions requires comparing the protein landscape of MN, their parental nuclei, and their surrounding cytoplasm, following distinct MN-generating genotoxic exposures.

Here, we used mass spectrometry-based proteomics to measure micronuclear protein composition from spontaneously-occurring MN (DMSO (vehicle)-treated cells) and MN generated from five exogenous stress conditions (3): Paclitaxel (Taxol), which produces MN containing whole lagging chromosomes; 10 Gy ionizing radiation (IR) inducing double-stranded breaks (DSB) among other varied DNA lesions (14); methylmethanesulfonate (MMS), a base- alkylating agent; hydroxyurea (HU), a replication stressor; and 5,6-dichloro-1-beta-D-ribofuranosylbenzimidazole (DRB), a transcription stressor. By comparing MN proteomes across distinct stress conditions, and by comparing MN proteomes to the nuclear and cytoplasmic proteomes from the same stress exposure, we provide a comprehensive assessment of the MN protein landscape. We find that the MN proteome is largely convergent across MN from all stress conditions examined here, indicating that global MN protein composition is not substantially influenced by upstream genotoxic exposure. MN are distinguishable from their parental nuclei by their significant lack of spliceosome machinery and DNA replication stress response components, and by their enrichment for a subset of the replisome. We demonstrate the utility of this resource by linking core MN proteome features to functional aspects of MN biology, establishing that that the loss of mRNA splicing machinery within transcribing MN contributes to intra-MN DNA damage, a known precursor to chromothripsis (7). This dataset describes a shared MN composition that is not altered by the upstream stress context, defining MN relative to their parental nuclear landscape and their surrounding cytoplasmic environment. Our observations offer new biological insight into the specific, vulnerable cellular processes that contribute to MN-dependent carcinogenic cascades.

## RESULTS

### The ionizing radiation (IR)-induced MN proteome recapitulates known MN features

We performed a first assessment of the MN proteome using 10 Gy IR-exposed MCF10A cells, whose MN features and downstream consequences have been previously characterized (3,5). Cells were allowed to expand for three days following IR exposure, at which time the MN and primary nuclei were separated by sucrose density gradients according to previously published protocols and subjected to mass spectrometry-based proteomic profiling (3,5,15) **(Fig. 1A)**. Irradiation, cell fractionation, and mass spectrometry runs were performed in triplicate for both MN and nuclear fractions, identifying 4,600 proteins in these six samples **(Table S1)**. For each detected protein, we compared the values measured in the MN fraction to the primary nuclei, to calculate both the log_2_ fold-change (FC) and the statistical significance (the *q-*value). A protein was considered significantly enriched or depleted in MN if it had both a log_2_ FC value greater than 1 (more abundant in MN than nuclei) or less than -1 (less abundant in MN than nuclei), and a *q*-value less than 0.05. Compared to their primary nuclei, IR MN from MCF10A cells are expected to carry high levels of the protein cGAS, and low levels of transcription machinery (3); they are also known to possess low levels of chromatin-remodeling proteins compared to their primary nuclei (1,16). These patterns were reflected in our measured IR MN proteome (**Fig. 1B**). MN envelopes have been shown to harbor similar or greater levels of core nuclear envelope proteins compared to their primary nuclei, including BAF and lamin A/C, while non-core envelope proteins such as lamin B and the nuclear pore complex components tend to be lacking in MN (2,17). This, too, was observed in our proteomes (**Fig. 1C**) and confirmed by immunofluorescent (IF) staining **(Fig. S1A-C)**. We next compared the IR MN proteome from MCF10A to that of a second human cell line, HeLa-S3, identifying 4,118 proteins **(Table S2)**. We found that a substantial fraction of the detected proteins were shared between MCF10A and HeLa IR MN (3,125 proteins, **Fig. S1D)**, and that among those shared proteins, there was a weak correlation in their average log_2_ FC values when compared to primary nuclei **(Fig. S1E)**. Where the absolute log_2_ FC value differed between HeLa and MCF10A, we found that the magnitude of the change between MN and nuclei is largely consistent **(Fig. S1E-F)**. One notable exception is cGAS, which is enriched in MCF10A but not in HeLa MN over primary nuclei **(Fig. S1F)**. However, this was expected due to the significantly higher levels of nuclear cGAS in HeLa cells, which alters the measured ratio of MN-to-nuclear cGAS abundance **(Fig. S1G)** (3). We conclude that the measured MN proteomes have recapitulated previously characterized MN features, and identified a largely consistent protein landscape in MN from two distinct human cell types induced by the same genotoxic exposure.

**Figure 1.**
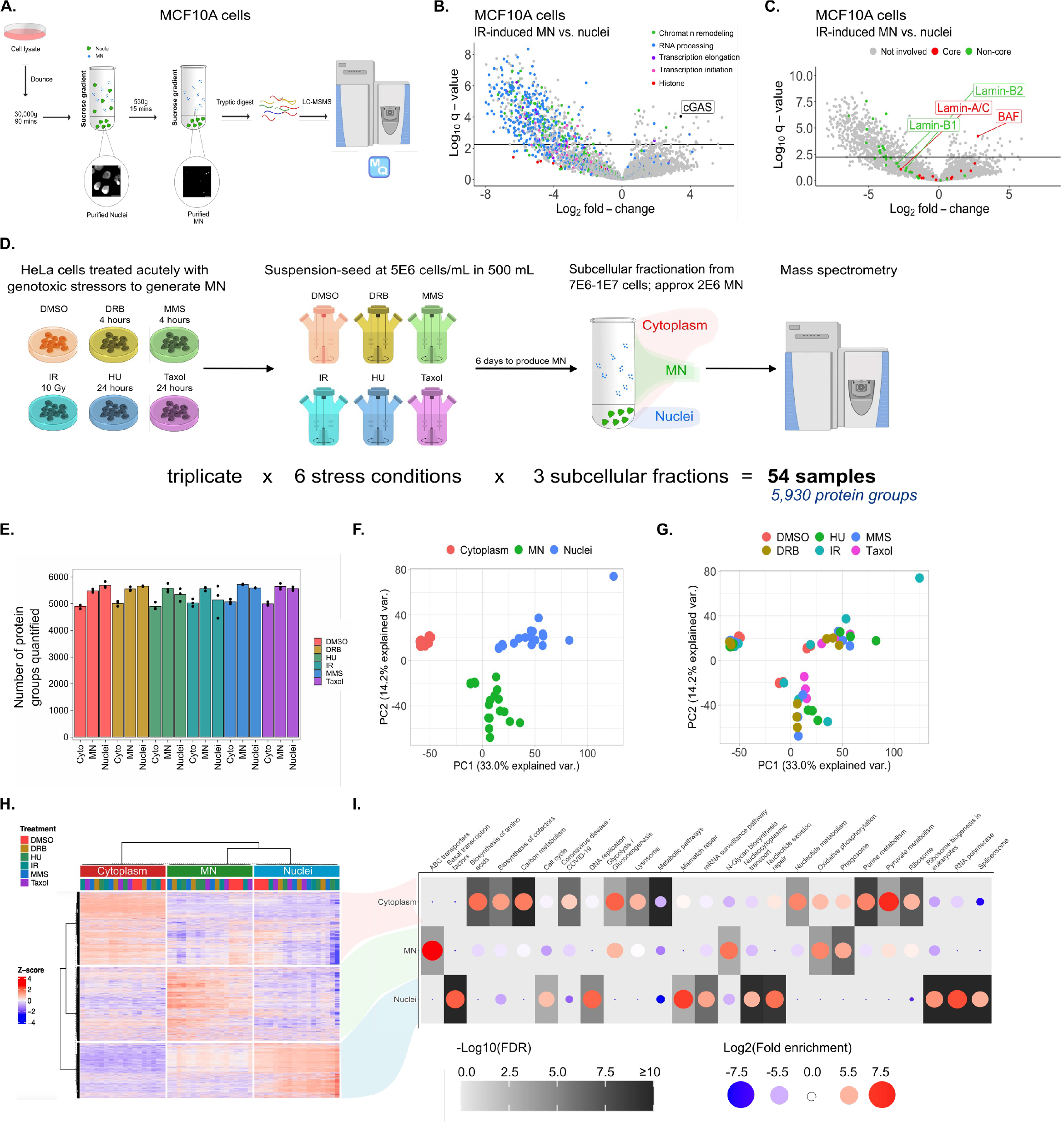
The micronuclear proteome is distinct from nuclei and cytoplasm and independent of inciting stress. **(A)** Workflow schematic for subcellular fractionation and whole proteomics in MCF10A. Cells are irradiated (10 Gy) or left untreated, in triplicate, and harvested 6 days later. Each sample is subjected to tryptic lysing, and peptides are quantified by liquid chromatography mass spectrometry (LC-MS) and MaxQuant software (MQ). Scale bar on representative micrographs of purified fractions = 20 μm. **(B-C)** Volcano plots displaying the log_2_ fold-change (FC) in proteins detected by MS, in 10 Gy IR MN compared to nuclei in MCF10A. Log_2_ FC for each protein (positive numbers indicating the protein is more abundant in MN) are plotted against the log_10_-transformed Benjamini-Hochberg (BH)-corrected t-test *p*-value (the *q-*value). Each point represents the mean log_2_ FC measurement for a single protein across three biological replicates. Horizontal black line indicates the cutoff for statistical significance using α = 0.05. (B) highlights chromatin and transcription proteins, (C) highlights core and non- core nuclear envelope proteins, as annotated by Gene Ontology. **(D)** Workflow schematic for treatment, fractionation, and proteome MS in HeLa-S3. Cells are grown in suspension, treated with each agent in our stress panel, and harvested 6 days later. Downstream processing as in (A). **(E)** The number of protein groups detected in each of the 54 HeLa samples by MS. **(F-G)** Principal component plots displaying the variance among the 54 HeLa samples based on MS intensity measurements. (F) is coloured by subcellular fractionation, (G) is coloured by genotoxic exposure. **(H)** Consensus clustering using MS intensity measurements, grouping samples by similarity. Each row represents a single protein, each column a single sample. Intensity measurements are presented as Z-scores, using the mean measurement for each protein across all 54 HeLa samples. **(I)** Result of KEGG gene ontology query, where the query list in is the proteins from each of the indicated clusters in (H) and the ∼6,000 proteins detected across all 54 samples is the background list. Only the top 25 terms (by *p-*value) are shown.

### The micronuclear proteome is distinct from nuclei and cytoplasm and independent of inciting stress

We aimed to understand two major features of the MN protein landscape. First, the distinctiveness of the MN proteome, relative to parental nuclei and resident cytoplasm; second, the degree to which the MN-generating genotoxic condition influences the global MN proteome, given prior work affirming that the recruitment of a small number of individual proteins is consistently affected by stress context (3,4). To address these questions, we expanded from IR to include DMSO, Taxol, MMS, HU, and DRB in our panel of MN-generating treatments. Due to the lower numbers of MN produced by some of these agents, large quantities of MN could not be obtained from MCF10A cells for mass spectrometry within reasonable bounds of starting material (3). We isolated cytoplasmic, MN, and nuclear fractions from 500 mL suspension cultures of HeLa-S3 cells treated with each of the six conditions in our panel, in triplicate **(Fig. 1D)**. Mass spectrometric assessments of all 54 samples detected in total 7,031 individual proteins **(Fig. 1E, Table S3)**. Using principal component analysis (PCA), we found that the MN proteome was clearly distinguishable from the nuclear and cytoplasmic proteomes, irrespective of the specific genotoxic stress condition used **(Fig. 1F-G)**. The proteins that are relatively high-abundance in the MN fractions were nearly evenly split between cytoplasmic and nuclear resident proteins, annotated by the Human Protein Atlas **(Fig. S2A)**. Within each of the cytoplasmic, nuclear, or MN sample clusters, there was no clear separation of samples by inciting stress **(Fig. 1G)**. Inciting stress-dependent MN recruitment of individual proteins such as cGAS has been previously reported (3,4) and was recapitulated in our proteome measurements **(Fig. S2B)**. However, the differences in specific protein abundance among MN were dwarfed by the distinctions between MN and nuclei or MN and cytoplasm **(Fig. 1G)**. Stress-specific comparisons of MN to nuclear or cytoplasmic protein abundance revealed several hundred proteins with significant differences in abundance, and the identity of these proteins is shared **(Fig. S2C-F)**. These observations confirm that MN are definable subcellular structures, accumulating a biased set of proteins rather than directly overlapping with either their parental nuclei or resident cytoplasmic compartment. MN consistently establish a core protein landscape that is only subtly affected by upstream stress context **(Fig. 1F-G)**.

We sought to identify and characterize the sets of proteins that define MN. Using *k*- means clustering to group all detected proteins by abundance, three broad groups emerged, representing proteins that are highest-abundance in either the cytoplasmic, MN, or nuclear fractions **(Fig. 1H)**. Considering the dataset in terms of MN, these are proteins that MN fail to recruit from their surrounding cytoplasm **(Fig. 1H top)**, fail to bring with them from their parental nuclei **(Fig. 1H bottom)**, or specifically recruit from either **(Fig. 1H middle)**. We characterized these groups by function using g:Profiler and the KEGG gene ontology dataset **(Fig. 1I)** (18). For a functional view of the dataset, we overlaid these three protein clusters onto the STRING protein-protein interaction network, highlighting the top ten (by *p*-value) functionally enriched pathways from KEGG **(Fig. 2A-B)** (19). Compared to their surrounding cytoplasm, MN are significantly depleted for ribosomal components, proteins involved in macromolecule metabolism, and the proteins of cytoplasmic-resident organelles including the lysosome **(Fig. 1H-I top, Fig. 2B-C)**. Proteins that are enriched in only MN compared to nuclear and cytoplasmic fractions are associated with the phagosome, as previously reported (20), and with oxidative phosphorylation **(Fig. 1H-I middle, Fig. 2B, Fig. 2D)**. Finally, relative to primary nuclei, MN fail to amass proteins involved in core nuclear processes including the DNA damage response (DDR), DNA replication, mRNA processing, and mRNA splicing **(Fig. 1H-I bottom, Fig. 2B, Fig. 2E-G)**. These results define the MN protein landscape by the presence and absence of functional classes of proteins, consistent with the concept of MN as distinct organelles.

**Figure 2.**
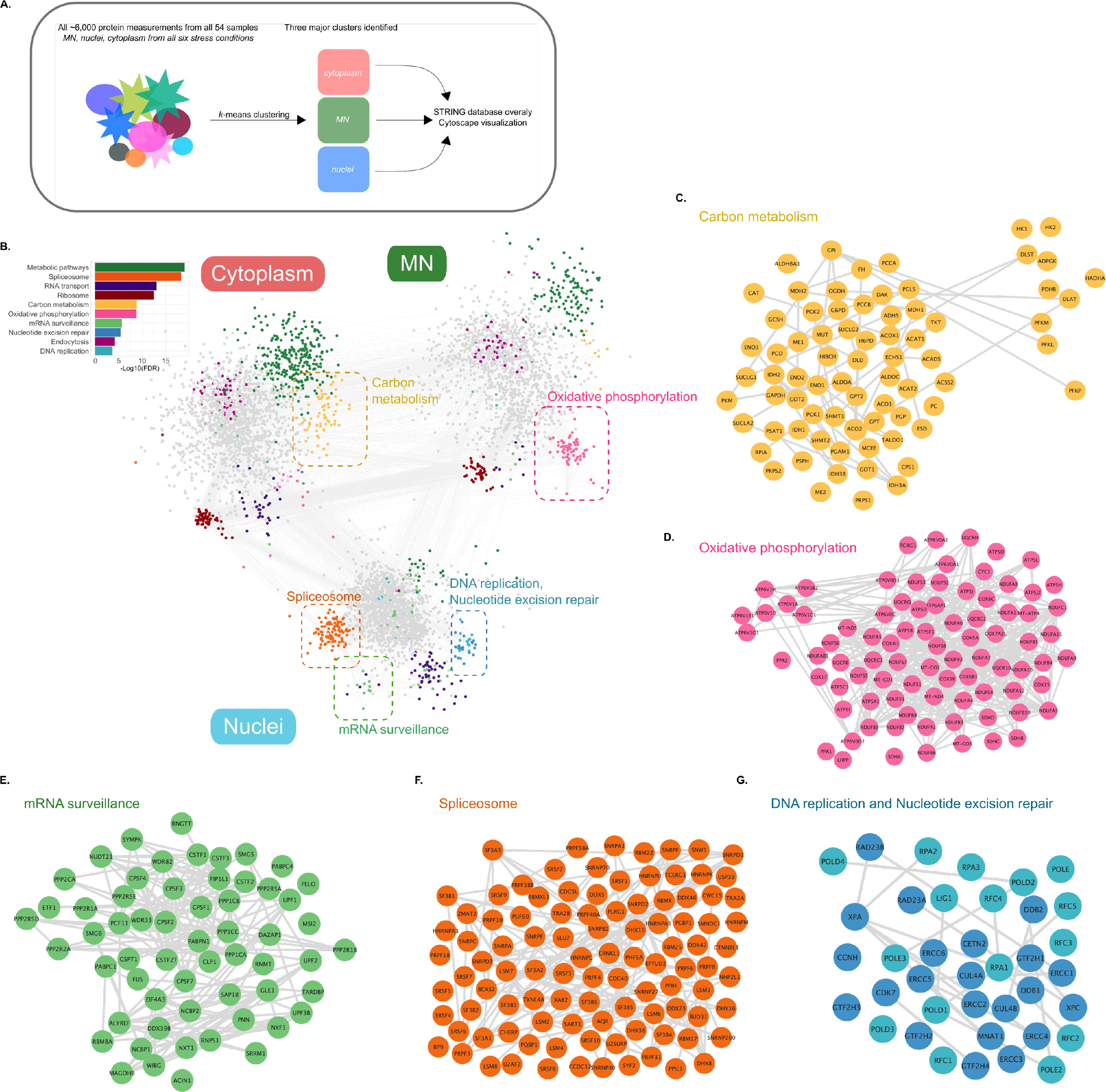
The functional protein landscape of MN, nuclei, and cytoplasm. **(A)** Conceptual depiction of the processing upstream of the functional network displayed in (B). **(B)** Functional interaction network for STRING protein clusters significantly high abundance in one of cytoplasmic, nuclear, or MN fractions (as in Fig. 1H). The ten subnetworks with the lowest false discovery rate (FDR)-corrected *p*-values are coloured and named based on KEGG queries of their protein constituents. Statistics of overrepresentation for each of these named subnetworks is displayed as a bar chart in the upper left part of the plot, with the log_10_-transformed *p*-value from Fisher’s exact test displayed on the x-axis, using an FDR cutoff of 5%. The constituents of the coloured subnetworks were used as the query list, and the ∼6,000 proteins in the full dataset were used as the background list. **(C-G)** Enlarged view of the carbon metabolism (B), oxidative phosphorylation (C), mRNA surveillance (D), spliceosome (E), and DNA replication and nucleotide excision repair (F) subnetworks presented in (A). For all networks, only edges with scores greater than 0.7 are shown.

### The micronuclear proteome mirrors the proteome of mitotic chromatin

We considered biological factors that could be responsible for the observed MN-specific protein profile, relative to surrounding cytoplasm and parental nuclei. MN are, fundamentally, fragments of double-stranded DNA or whole chromosomes that are not metabolically active, explaining their failure to attract proteins associated with carbon metabolism or macromolecule processing once they are resident in the cytoplasm **(Fig. 1H-I top, Fig. 2B-C)**. Associations between MN and the phagosome (**Fig. 1H-I middle)** and lack of interaction between cytosolic DNA and the ribosome **(Fig. 1H-I top)** have been previously observed (12,20). The reasons for MN- enrichment of proteins involved in oxidative phosphorylation are less clear, as this is a process typically restricted to the mitochondria. Initially, we considered that mitochondria could co-sediment during MN isolation (21). However, when we remove all proteins tagged as mitochondrial residents by Gene Ontology from our dataset (638 proteins in total; nearly 10% of all proteins identified in this dataset, **Table S4**), MN samples remain uniquely clustered, indicating that mitochondrial proteins do not drive the distinctiveness of the MN proteome **(Fig. S2G)**. The enrichment for oxidative phosphorylation in MN even when compared to the cytoplasmic fraction also supports true enrichment of the proteins, rather than the mitochondria themselves, since the cytoplasmic fraction is not depleted of mitochondria prior to preparation for mass spectrometry **(Fig. 1H-I top)**. Associations between mitochondrial proteins and MN DNA remain an area of active study. cGAS, primarily considered a viral and MN DNA-binding protein, also recognizes mitochondria (22–24), but to our knowledge no examination of whether mitochondrial DNA-binding proteins can bind MN DNA has been performed. Of the twenty-six proteins considered by Gene Ontology to be associated with mitochondrial DNA, eleven are relatively enriched in the MN fractions, among them uniquely mitochondrial DNA-binding proteins including POLG and SSBP1 **(Fig. S2H)**. Immunofluorescent staining confirmed the frequent presence of these proteins in MN from all tested stressors **(Fig. S2I)**. In contrast, MGME1, a mitochondrial nuclease, and CHCHD4, which aids in the formation of disulphide bonds during cellular respiration, were found enriched in the cytoplasmic fraction of our proteome dataset **(Fig. S2H)** and in relatively few MN by IF staining **(Fig. S2I)**. These observations suggest that certain cytoplasmic proteins, including phagosome components or mitochondrial DNA-binding factors, are recognizing MN DNA once in the cytoplasm while other cytoplasmic residents, such as those associated with macromolecule metabolism or protein translation, are excluded.

We next considered the proteins that are significantly depleted in MN compared to primary nuclei. When they do not contain whole chromosomes MN are, by definition, regions of the genome where a DSB has occurred and DNA damage repair has failed. The appearance of *homologous recombination* as a significantly depleted ontology term in MN is consistent with this model **(Table S5)**. However, MN are also significantly lacking in proteins associated with terms such as *DNA replication, nucleotide excision repair, mismatch repair, RNA processing, regulation of mRNA stability* and, the most significantly altered KEGG pathway, *spliceosome*, none of which are unambiguous routes to unresolved DSB **(Fig. 1I bottom, Table S5)**. Instead, these pathways are consistent with those previously shown to be depleted from mitotic chromatin (25). Comparing MN proteomes to the proteomes of G1-, S-, and M-phase chromatin using a publicly available dataset (25), our MN proteome was most similar to that of mitotic chromatin **(Fig. S3A-C)**. Both mitotic chromatin and MN were relatively depleted for spliceosome machinery and small-molecule metabolism factors, and relatively enriched for oxidative phosphorylation proteins, within their respective datasets **(Fig. S3D-E)** (25). Together, these measurements suggest that elements of the core MN proteome, shared by all MN examined here, could be acquired from several sources: exclusion of cytoplasmic residents associated with metabolism or translation, due to a fundamental MN composition of DNA; recruitment of DNA-binding cytoplasmic residents, including those from the mitochondria; and the large-scale changes to mitotic chromosomes that occur at the time of MN sequestration, stripping whole classes of nuclear proteins which MN cannot easily recover (2,26).

### Protein distribution is subtly but distinctly shifted when MN are generated by specific stress exposures

Proteomic profiling has revealed that the protein landscape of stress-induced MN is largely convergent **(Fig. 1F-H)**. However, significant differences in the recruitment of individual, highly consequential proteins to MN generated by a panel of stressors has been previously observed by several groups, including ours (3,4,27). These include the pattern recognition receptor (PRR) cGAS, transcription machinery, and certain DDR proteins (1,3,4,7,28). While we have observed a shared protein profile in bulk among MN from distinct stress conditions, we asked whether the distribution of only a small number of proteins within MN could betray the specific upstream genotoxic context, with signatures identifiable by proteomic profiling. Consistent with this concept, while known functional groups of proteins are enriched or depleted in all MN compared to all nuclei in our panel **(Fig. 3A)**, there is subtle heterogeneity among MN by stress condition, with statistically significant differences in protein abundance **(Fig. 3B-D)**. We first asked whether individual proteins, present or absent in MN from each stress context, could act as biomarkers identifying MN as being the product of a particular genotoxic lesion. We set the cutoff for an individual protein as an MN biomarker candidate if it was, first, enriched only in the MN fractions compared to nuclei and cytoplasm, and second, more than three times as abundant in MN from that stressor compared to MN from all five of the other stress conditions (a log_2_ FC cutoff of 1.58). Only fifteen proteins out of the total dataset of nearly 6,000 passed these filters **(Fig. S4A)**. These proteins were not significantly associated with any functional pathway by GO, and had no significant interactions within the STRING database. Individual factors may be used to identify MN from specific stress contexts and can be profiled by IF staining **(Fig. S4B)**, but this analysis suggests that the presence of those proteins in MN has no specific relationship to the cellular response to that genotoxin.

**Figure 3.**
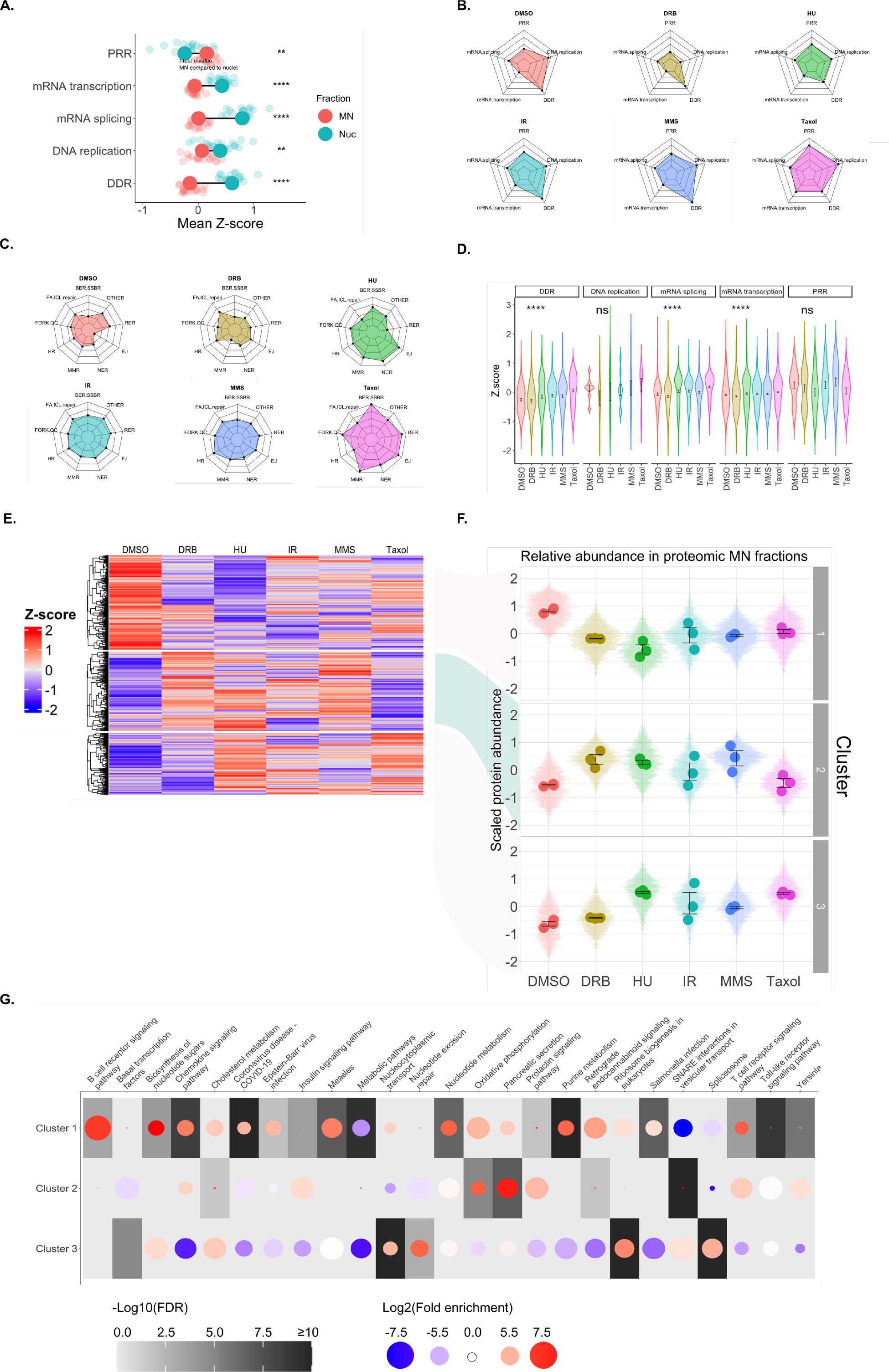
Protein distribution is subtly but distinctly shifted when MN are generated from specific stress exposures. **(A)** Z-score intensity measurements by mass spectrometry (MS) for proteins from each of the indicated categories, in MN samples compared to nuclei samples. Each translucent point represents the mean Z-score for a single protein, across three biological replicates, from a single genotoxic stress condition. Each large solid point represents the mean scaled intensity measurement for all proteins in that category, measured in the MN or nuclear samples. **(B-C)** Radar plots displaying the scaled intensity measurements by MS for proteins from each of the indicated categories, in MN samples from each of the stress conditions in our panel. The outer edge of the plot represents the largest intensity measurement across all samples, and the center of the plot represents the smallest intensity measurement. **(D)** Z-score intensity measurements for proteins from each of the indicated categories, in MN samples from each of the stress conditions in our panel. Violin plots show the distribution of measurements for proteins in that category. Statistical comparisons by one-way ANOVA. **(E)** Consensus clustering using MS intensity measurements, grouping samples by similarity. Each row represents a single protein, each column the mean of three biological replicates for MN from the indicated stress condition. Intensity measurements are presented as Z-scores, using the mean measurement for each protein across all MN samples. **(F)** Mean Z-scores represented in (E). Violin plots show the distribution of Z-scores for proteins from the indicated stress condition in the indicated cluster. Large solid points represent the mean Z-score for all proteins in that cluster, from a single biological replicate of from stress-induced MN. **(G)** Results of KEGG gene ontology query, where the query list in is the proteins from each of the indicated clusters in (E) and the proteins found in all MN samples is the background list. Only the top 25 terms (by *p-*value) are shown. All error bars represent standard error of the mean, for three independent biological replicates. For all statistical comparisons, ns: *p* > 0.05, *: *p* <= 0.05, **: *p* <= 0.01, ***: *p* <= 0.001, ****: *p* <= 0.0001.

Rather than limiting our analysis to proteins that are significantly high-abundance in MN from only a single stress condition, we considered proteins with a consistent distribution among groups of stress-induced MN. One known example is cGAS, which is high-abundance in MN generated following exposure to IR or MMS, and low-abundance in MN that form following DMSO, HU, and DRB exposures, as measured both in our proteome samples **(Fig. S2B)** and in prior work (3,4). We asked whether our dataset could identify other proteins like cGAS, with a consistent distribution among groups of stress-induced MN. We performed *k*-means clustering on only the MN samples from each of the six exposure conditions, grouping proteins by similarity in their relative abundance **(Fig. 3E)**. Three clusters of proteins emerged: proteins that are high-abundance in only DMSO MN **(Fig. 3E-F top**, “Cluster 1”); relatively high-abundance in DRB, HU, IR, and MMS MN while low-abundance in DMSO and Taxol MN **(Fig. 3E-F middle**, “Cluster 2”); and high-abundance in HU, IR, MMS, and Taxol MN while low-abundance in DMSO and DRB **(Fig. 3E-F bottom**, “Cluster 3”). Cluster 1 proteins are relatively high-abundance in only DMSO MN, potentially indicating a distinction between spontaneously-occurring MN and those induced by exogenous genotoxins. This distinction was also observed when comparing DMSO nuclei to all others suggesting these MN differences arise from the overall changes in protein abundance within the cell after genotoxic stress **(Fig. S4C-D)**. Cluster 2 proteins, which include cGAS, tended to be associated with oxidative phosphorylation **(Fig. 3F-I)**. Cluster 3, low-abundance in DMSO- and DRB-induced MN only, contains proteins significantly associated with the spliceosome **(Fig. 3G)**. Taken together, our observations indicate that while the core MN protein landscape is convergent and definable **(Fig. 1F-I, Fig. 2B)**, the specific MN-generating stress condition induces subtle redistributions in nuclear and cytoplasmic protein abundance, with functional protein groups distributing together **(Fig. 3)**.

### Micronuclei are characterized by spliceosome deficiency and replisome enrichment

We have observed that, compared to primary nuclei, all MN from our panel are significantly depleted for proteins involved in DDR, transcription, chromatin modification, and DNA replication, in part attributable to removal of these proteins from mitotic chromatin prior to MN formation **(Fig. 2, Fig. S3)**. However, prior work has established that some MN are able to recover a subset of nuclear proteins, and MN do engage in abortive or delayed versions of those core nuclear processes, which risks DNA degradation and chromothriptic rearrangement of the MN-sequestered chromosome (1,3,7–9,13). One mechanism contributing to this degradation, incomplete base excision repair, has been described (7), but the full landscape of nuclear machinery that drives chromothriptic DNA damage within MN is not known. Furthermore, the analysis presented above suggests subtle but significant redistribution of proteins involved in DDR, DNA replication, and transcription within MN by stress condition **(Fig. 3)**. Given that the DNA degradation upstream of chromothripsis depends on incomplete DNA replication, transcription, and DDR within the MN, these redistributions could influence the risk of chromothripsis after specific stress exposures (7,13). We asked whether the MN proteome could identify functional groups of nuclear proteins that contribute to MN DNA degradation after genotoxic stress. To focus on nuclear factors that are part of the core MN protein landscape, we pooled our MN protein measurements across the six genotoxic conditions in our panel, and performed differential expression analysis for mean protein levels comparing to the pooled nuclear samples. We found 1,489 significantly depleted and 1,466 significantly enriched proteins in MN compared to primary nuclei using this pooled approach **(Fig. S4E, Table S6)**. Next, we computed the Pearson correlation coefficient (PCC) for the mean detected protein level for each pair of proteins in the enriched list of 1,466 or the depleted list of 1,489, discarding any pair that had a PCC less than 0.7 **(Table S7)**. This step allows us to focus on a set of proteins whose presence or absence in MN is strongly intercorrelated, increasing the chance of identifying a functional interaction (29). Finally, we overlaid our list of candidate proteins onto the STRING protein-protein interaction database (19). On the MN-depleted side, 1,172 proteins corresponded to STRING identifiers. Of those, 158 were “singletons,” not known to functionally interact with any other protein in the list, and another 6 proteins interacted with only one other protein. This left us with a highly interconnected network of 1,008 terms, where nodes are proteins that are significantly depleted in all MN compared to nuclei and edges represent both a functional interaction recorded in STRING and a PCC of at least 0.7 within our MN proteomes dataset **(Fig. 4A-B)**. Functional enrichment analysis revealed visible submodules within this network, significantly associated with the spliceosome, DNA damage repair sub-pathways, DNA replication, and RNA polymerase subunits **(Fig. 4B-D)**. In particular, the DDR submodule significantly under-represented in all MN is an ATR-governed network **(Fig. 4C)**. On the MN-enriched side, 1,196 proteins corresponded to STRING identifiers, with 5 singletons **(Fig. 4E)**. We found that MN are significantly enriched for a subnetwork of viral receptors and proteins associated with the innate immune system, consistent with published work **(Fig. 4E)** (3,5,6). MN are also enriched for proteins associated with M-phase, oxidative phosphorylation, and mitochondrial translation, recapitulating the significant overlap between our MN proteome dataset and the mitotic chromatin-associated proteome published in (25) **(Fig. 4E, Fig. S3D-E)**. Within MN-enriched proteins, a small submodule of replisome components emerged including the RFC and MCM family proteins, DNA polymerase subunits, and single-stranded DNA-binding proteins including RPA1 and FEN1 **(Fig. 4F)** (30). Using functional interaction network analysis of a significantly correlated fraction of MN-enriched and -depleted nuclear proteins, we have identified visible submodules associated with DNA replication, ATR-dependent DDR, M-phase chromatin, and the mRNA spliceosome.

**Figure 4.**
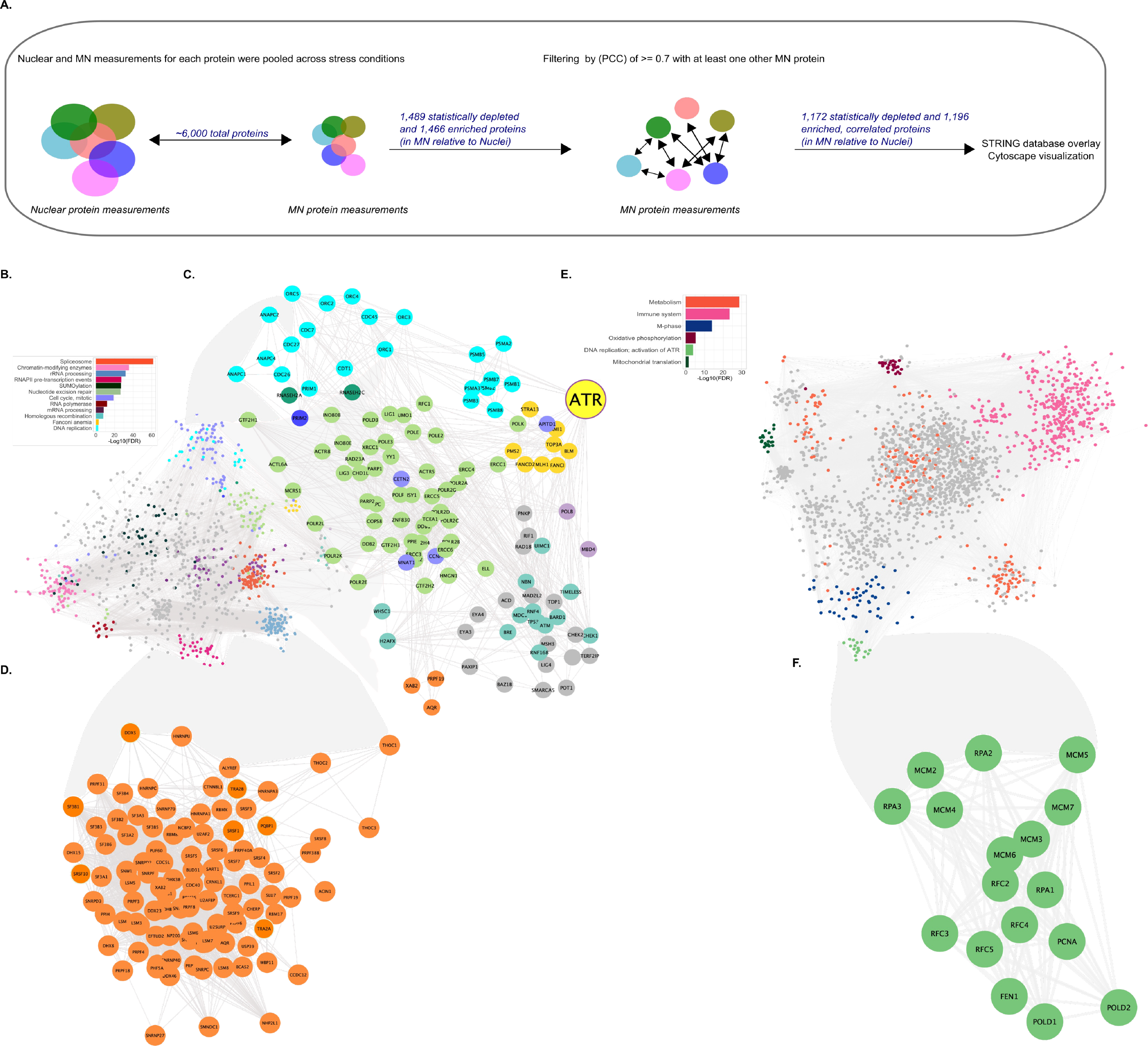
Micronuclei are characterized by spliceosome deficiency and replisome enrichment. **(A)** Conceptual depiction of the processing upstream of the functional network displayed in (B). In step (1), each pooled protein measurement was compared between MN and Nuclei by Benjamini-Hochberg (BH)-corrected t-test. In step (2), the original MN protein abundances of each significantly enriched or depleted protein were compared to each of the other enriched or depleted proteins. Only those proteins that had a Pearson correlation coefficient (PCC) of >= 0.7 with at least one other MN protein passed the filter. **(B)** Functional interaction network for proteins significantly depleted in all MN compared to all primary nuclei. Significantly depleted proteins whose measured abundance in MN correlates with at least one other protein with a Pearson correlation coefficient of 0.7 or greater were queried against the STRING protein-protein interaction network database. Visually identified subnetworks were coloured and named based on KEGG queries of their protein constituents. Statistics of overrepresentation for each of these named subnetworks is displayed as a bar chart in the upper left part of the plot, with the log_10_-transformed *p*-value from Fisher’s exact test is displayed on the x-axis, using a false discovery rate (FDR) cutoff of 5%. The constituents of the coloured subnetworks were used as the query list, and the 1,172 depleted proteins in the full network were used as the background list. **(C)** Enlarged view of the DNA replication and DDR subnetworks. ATR is indicated by a larger node with a purple border. **(D)** Enlarged view of the spliceosome subnetwork. **(E)** The same analysis as (A), but using the proteins found significantly enriched in all MN compared to all nuclei. **(F)** Enlarged view of the DNA replication; activation of ATR signalling subnetwork.

### Absence of spliceosome proteins in transcriptionally competent MN contributes to MN DNA degradation

Given the significant depletion for spliceosome components in MN **(Fig. 4B, Fig. 4D)**, and the identification of spliceosome machinery among those groups of MN proteins that may depend on the inciting stress condition **(Fig. 3E-G, bottom row**), we asked whether aberrant splicing could contribute to the prior observation that MN DNA degradation, a precursor to chromothripsis, is transcription-dependent (7). We first aimed to establish whether splicing machinery, like transcription machinery, is heterogeneously distributed across MN (3). We used IF to score the fraction of MN carrying the proteins SRSF1 and SF3B1, two core spliceosome components that are significantly low-abundance in MN compared to nuclear fractions across all our measured stress conditions **(Fig. 5A, Table S6)**. A subset of MN from all treatments recruited SRSF1 and SF3B1, at frequencies depending on the treatment and commensurate with the level of active transcription marked by 5-ethynyl uridine (EU) incorporation **(Fig. 5A)** (3,4). Thus, while mRNA splicing factors are depleted in bulk in the MN proteome, this reduction is not uniform across all MN, and its magnitude depends on the specific stress condition used.

**Figure 5.**
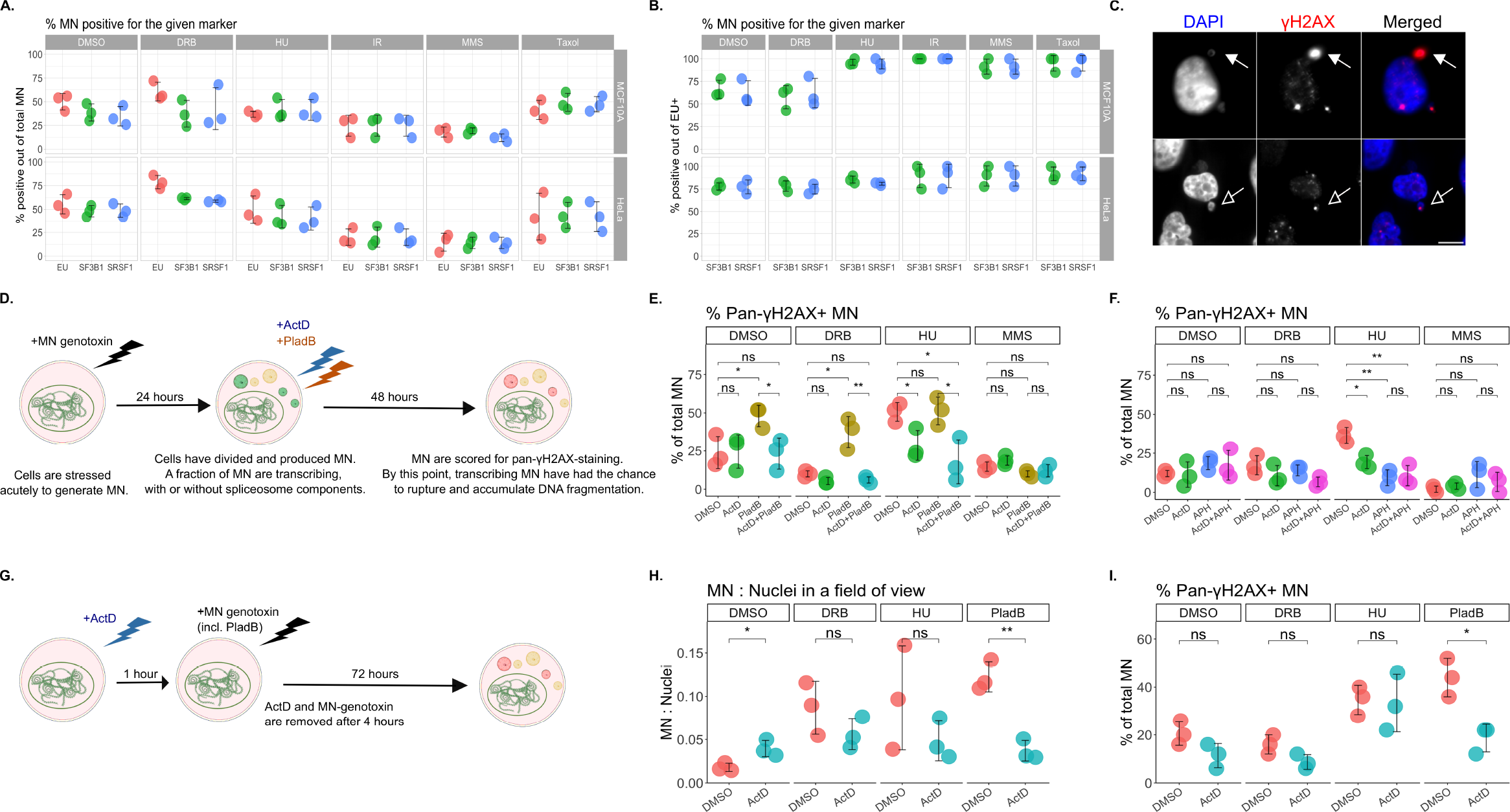
Absence of spliceosome proteins in transcriptionally competent MN contributes to MN DNA degradation. **(A-B)** Percent EU+, SF3B1+, or SRSF1+ MN by immunofluorescence (IF), 72 hours following each of the indicated acute stress conditions in MCF10A (top) and HeLa (bottom). (A) presents the percentage of IF marker-positive MN as a fraction of total MN, while (B) is as a fraction of the EU+ MN (i.e. co-localization with SF3B1 or SRSF1). **(C)** Representative pan-γH2AX+ MN (filled arrow, top), and foci-γH2AX+ MN (empty arrow, bottom) in MCF10A. Scale bar = 10 μm. Image representative of 3 experiments. **(D)** Experimental timeline for perturbing splicing and transcription within MN prior to MN DNA fragmentation. **(E-F)** Percent pan-γH2AX+ MN in MCF10A by IF when perturbing splicing with pladienolide B (PladB), transcription with actinomycin D (ActD), or replication with aphidicolin (APH) using the experimental timeline in (D). **(G)** Experimental timeline for perturbing splicing and transcription in nuclei during MN-generative acute genotoxic exposure. **(H)** Number of MN per nuclei present in a microscopy field of view (FOV), or **(I)** Percent pan-γH2AX+ MN by IF in MCF10A, following the experimental timeline in (G). All statistical comparisons performed using a two-sided Student’s t-test. NS: p=1, ns: *p* > 0.05, *: *p* <= 0.05, **: *p* <= 0.01, ***: *p* <= 0.001, ****: *p* <= 0.0001. For (H), each point represents the mean of at least 5 FOVs, for a total of three independent biological replicates. All other individual data points presented for IF scoring of MN represent the mean percentage of MN that were positive for the indicated marker, from each biological replicate out of 50 total MN per replicate. All error bars represent standard error of the mean, for three independent biological replicates.

SRSF1+ or SF3B1+ MN were preferentially localized to actively transcribing (EU+) MN, but we also observed a subset of MN that were engaged in transcription without SF3B1 or SRSF1 **(Fig. 5B)**. We asked whether MN that are transcriptionally competent, but unable to splice mRNA, are at particular risk for intra-MN DNA damage. MN DNA fragmentation can be scored by IF staining for γH2AX, when present as a bright, diffuse pattern spanning the entire MN **(Fig. 5C top)**; small γH2AX foci represent individual DSB within MN, and are not scored as chromosome fragmentation **(Fig. 5C bottom)** (7,28). We exposed MCF10A cells to our panel of acute MN-generative genotoxins, allowing 24 hours to produce MN. At this point, the majority of MN have not ruptured their MN envelope, and are capable of active transcription (1,3). We then added 20 nM pladienolide B (PladB; a pharmacological inhibitor of SF3B1) to inhibit splicing, and/or 0.01 μg/mL actinomycin D (ActD) to prevent transcription, and allowed cells to progress for a further 48 hours. During these 48 hours, the MN generated from this genotoxic panel experience maximum envelope rupture (3), and are vulnerable to cytoplasmic processing and MN DNA damage (7) **(Fig. 5D)**. Using this experimental setup, we observed stress-specific patterns of MN DNA damage. At baseline, MN from DMSO, DRB, or MMS exposure exhibit low levels of pan-γH2AX staining, while half of all MN from HU show γH2AX patterns consistent with MN DNA degradation **(Fig. 5E)**. DNA degradation in HU MN was significantly reduced by inhibiting transcription with ActD; in contrast, baseline DNA damage in DMSO-, DRB-, and MMS-induced MN does not appear to be transcription-dependent **(Fig. 5E, Fig. S4F)**. In the MMS-treated condition, this is likely the result of low overall transcriptional activity in MN **(Fig. 5A)**. DMSO- and DRB-induced MN, however, are transcriptionally competent, and frequently lack SF3B1 or SRSF1 **(Fig. 5A-B)**. We found that inhibition of MN splicing using PladB significantly increased pan-γH2AX+ MN following DMSO and DRB only **(Fig. 5E)**. This increase was dependent on transcription, abrogated by concomitant ActD treatment, where PladB treatment alone does not significantly alter the incidence of transcribing MN **(Fig. 5E, Fig. S4F)**. Thus, under conditions where a large fraction of MN are transcriptionally competent (such as DMSO and DRB exposure), cells appear to be at particular risk for the MN DNA fragmentation and chromothripsis associated with abortive intra-MN transcription, especially when intra-MN transcription occurs in the absence of a functional spliceosome **(Fig. 5B-E)** (7).

Notably, the majority of HU-induced MN are pan-γH2AX+ at baseline, partially driven by transcription but with no clear role for splicing defects **(Fig. 5E)**. Aberrant intra-MN replication has also been linked to MN DNA damage (7,8,13), and altered distribution of replisome and replication stress response components were identified in our functional network analysis of MN **(Fig. 4C, Fig. 4F)**. We asked whether, in the HU-exposed condition, MN DNA replication was a significant contributor to baseline MN DNA damage. When we paused MN DNA replication with 3 μM aphidicolin (APH) using the same experimental timeline as used for splicing inhibition **(Fig. 5D)**, HU-induced MN were significantly reduced in their pan-γH2AX staining **(Fig. 5F)**. No reduction was observed in DMSO-, DRB-, or MMS-induced MN **(Fig. 5F)**. Preventing MN transcription alongside MN replication had no additive effect **(Fig. 5F)**. These observations suggest that despite convergence in the identity of their nuclear resident proteins **(Fig. 4)**, the functional consequences of that shared protein landscape depend on the genotoxic origins of MN. This prompted us to ask whether MN originating from nuclear splicing or transcription defects, rather than MN that have splicing or transcription defects imposed after their formation, would exhibit unique propensity for MN DNA degradation. We again scored pan-γH2AX+ MN following exposure to our panel, but included an acute high-dose exposure to PladB (500 nM for four hours) among the stress conditions, and induced MN with or without nuclear transcriptional capacity using concomitant treatment with ActD **(Fig. 5G)**. Significant MN formation was induced with exposure to 500 nM PladB, but this was dependent on nuclear transcriptional activity during the treatment window **(Fig. 5H)**. PladB-induced MN, like HU-induced MN, were pan-γH2AX+ in the majority by 72 hours post-exposure **(Fig. 5I)**. The induction of MN while nuclear transcription was paused with ActD significantly reduced % pan- γH2AX+ MN induced by PladB alone **(Fig. 5I)**. We conclude from these results that while specific genotoxic stressors play relatively minor roles in dictating the core protein landscape that define MN **(Fig. 1F-H, Fig. 3)**, they confer specific, exploitable vulnerabilities for MN DNA degradation, a known precursor to chromothripsis following MN transcription or replication **(Fig. 5)** (7). Examination of the micronuclear proteome has identified a previously unknown role for splicing machinery in MN generation and activity, emphasizing the functional utility of mapping the MN protein landscape.

## DISCUSSION

Micronuclei are byproducts of mitotic progression despite unresolved DNA damage, arising in both stressed normal cells and chromosomally unstable cancer cells (3,5–7,9). Once formed, MN can have profound consequences for the cell and its daughters, notably through viral pattern recognition receptor recruitment (3,5,6) or the introduction of chromothriptic gene rearrangements (7,9). These MN-driven consequences depend on their DNA and protein content, meaning that the comprehensive mapping of MN properties is crucial for expanding our understanding of cancer development and progression (2–4,7,8,11,12). Here, we have used mass spectrometry to profile and directly compare the proteomes of MN generated from a panel of distinct genotoxic stress conditions, and their matched nuclear and cytoplasmic fractions. This dataset allowed us to address two major, open questions in micronuclear biology: First, the relative influence of parental nuclei or surrounding cytoplasmic pools on MN protein composition; and second, the influence of the specific MN-generating genotoxic stress context on the MN protein landscape. Our analysis of MN from six stressors has revealed more convergence than divergence in their protein content **(Fig. 1F-H)**. Micronuclei, regardless of the specific stress context, are a meaningful subcellular compartment with a consistently measured proteome, defined by enrichment for certain replisome components and immune signalling proteins and lack of DDR, mRNA splicing, and macromolecule metabolism **(Fig. 2)**. The MN protein composition we have observed here appears to reflect several non-exclusive processes, beginning with processing of mitotic chromatin **(Fig. S3)**. Once formed and cytoplasm- sequestered, the MN protein landscape can be further shaped by their lack of association with metabolic factors and their recruitment of cytoplasmic DNA-binding proteins such as viral receptors and mitochondrial proteins **(Fig. 1H-I, Fig. 2B-D, Fig. S2H-I)**, as well as a whole-cell protein redistribution in response to stress **(Fig. S4C-D)**. This work establishes a core MN proteome, amassed from both parental nuclei and surrounding cytoplasm, and only subtly affected by the specific conditions of their formation **(Fig. 3)**.

The convergence in our MN proteomes, particularly on the absence of nuclear DDR factors and core nuclear activities including transcription and splicing, suggested to us that there may be common, vulnerable nuclear processes that dictate MN function under many stress contexts. MN DNA degradation, upstream of chromothripsis and marked by a pan-MN pattern of γH2AX has been previously linked to abortive MN transcription or replication **(Fig. 4D)** (7). Following our proteomic identification of mRNA splicing as a significantly, consistently depleted functional network within MN, we showed that shared, impaired splicing is another contributor to pathological MN DNA damage **(Fig. 5E)**. However, despite their overlap in functional enrichment or depletion, the conditions of MN formation do alter their protein content **(Fig. 3)**, and small differences in MN protein composition can have large functional impacts (3). We show here that within DMSO- and DRB-generated MN, the modestly lower abundance of splicing machinery within actively transcribing MN manifests as a transcription-dependent pan- γH2AX signature consistent with MN DNA fragmentation **(Fig. 5A-E)** (7). This dependency was not shared by MMS-induced MN, which largely do not transcribe, or HU-induced MN, where DNA degradation was driven by MN DNA replication rather than only splicing defects **(Fig. 5A- F)**. Thus, alongside a core MN proteome established irrespective of stress exposure, our work identifies stress-specific MN protein signatures, where variance in individual protein recruitment to MN can betray the specific circumstances of their formation and significantly affect MN-driven outcomes. The processes responsible for this stress specificity remain to be fully understood, but may stem from differences in local nuclear chromatin organization induced by different genotoxins, including alterations to histone modifications or propensity for unresolved ssDNA gaps (3,4). Datasets such as the one we have generated here, which comprehensively profile the role of stress exposure on the cytoplasmic, nuclear, and micronuclear protein landscape, will guide mechanistic studies of MN-generation following specific forms of genotoxic stress. More broadly, this work demonstrates that consistent distribution among MN from specific stress conditions is a quality shared by many proteins, expanding from the previously observed cGAS and transcriptional machinery (3).

In summary, our work represents the first direct comparison of MN protein content across discrete genotoxic exposures. The dataset that we have generated can be used to identify functionally relevant proteins and pathways involved in MN formation and consequential MN-driven cellular processes, representing a key resource for future work investigating the progression of genomic instability following genotoxic exposure. MN are the direct result of failure to resolve a wide variety of potential genotoxic stress conditions, and this resource can be used to identify common, MN-generating vulnerabilities within the stressed cell.

## ACKNOWLEDGEMENTS

This work in the Harding lab was supported by a Canadian Institutes for Health Research (CIHR) Project Grant (PJT165926), the National Science and Engineering Research Council of Canada (NSERC) Discovery Grant (RGPIN 2019 0481), the JP Bickell Foundation, the Princess Margaret Cancer Centre, the Princess Margaret Cancer Foundation and Ontario Ministry of Health. KMM has received support from the Ontario Graduate Scholarship, CIHR, and the University of Toronto Endowed Awards. This work was partially funded through a CIHR Project Grant (PJT 173487) to T.K. T.K. is supported through the Canadian Research Chair program.

## AUTHOR CONTRIBUTIONS

KMM and SMH designed the experiments and wrote the manuscript with input from all authors. All cellular experiments were completed by KMM. Mass spectrometry experimental design, sample preparation and data collection and processing were performed by SK and TK with input from KMM and SMH. Data analysis was conducted by KMM with input from SK. Final figure and table preparation was performed by KMM and SK.

## DATA AVAILABILITY

Mass spectrometry raw data have been deposited to the Mass Spectrometry Interactive Virtual Environment (MassIVE).

## METHODS

### Cell culture

MCF10A cells (ATCC cat #CRL-10317) were cultured in 1:1 mixture of F12:DMEM media supplemented with 5% horse serum (Wisent Bioproducts cat #098150), 20 ng/ml human EGF (Cedarlane Labs cat #AF-100-15), 0.5 mg/ml hydrocortisone (Sigma cat #H0888), 100 ng/ml cholera toxin (Sigma Aldrich cat #C8052) and 10 mg/ml recombinant human insulin (SAFC cat #91077C). HeLa-S3 cells (ATCC cat #CCL-2) were grown in DMEM supplemented with 10% FBS. All media was supplemented with 1% penicillin-streptomycin. All cell lines were authenticated by ATCC using short tandem repeat sequencing. For experiments involving drug treatments, drugs were added directly to the cell media. For our panel of six acute MN- generating genotoxic stress conditions (Fig. 1D), doses and exposure times were selected based on existing literature and preliminary experiments to induce DNA lesions and maximize MN formation, while minimizing cell death and cell cycle arrest within the time frame of the assay (3-6 days, depending on the experiment) (3). Doses and exposure times used for each agent are as follows: ionizing radiation (IR, Cs-137), 10 Gy (5); methylmethanesulfonate (MMS, Sigma cat #12995), 0.5 μM for 4 hours (31); paclitaxel (Taxol, Selleck Chemicals cat #S1150), 10 nM for 4 hours (32); hydroxyurea (HU, Sigma cat #148627), 2 mM for 24 hours (32,33); 5,6- dichloro-1-beta-ribo-furanosyl benzimidazole (DRB, Sigma cat #D19116), 100 μM for 4 hours (34). At the end of the designated exposure time, cells were washed in PBS, replaced with fresh media, and allowed to cycle freely for 72 hours to produce MN before proceeding to downstream analysis. For the experiments presented in Fig. 5E-F, MCF10A cells were exposed to the indicated acute stress condition from our six-condition panel, then allowed 24 hours to produce MN. Cells were then treated with 20 μM pladienolide B (PladB) to inhibit splicing for a further 48 hours, 0.01 μg/mL actinomycin D (ActD) to prevent transcription, or 3 μM aphidicolin (APH) to prevent replication. During these 48 hours, the MN from these stress conditions experience maximum envelope rupture (3) and are vulnerable to cytoplasmic processing and MN DNA damage as reported in (7). MN were then scored as described under the subheading *immunofluorescent miscroscopy*.

### Micronuclear, nuclear, and cytoplasmic subcellular fractionation

Performed as publishes and described in (3) with the following modification: Cytoplasmic fractions (supernatant) were collected immediately following the cell lysis and centrifugation step rather than discarded.

### Sample preparation for shotgun proteomics

Micronuclei and nuclei fractions were lysed by repeated freeze-thaw cycles in lysis buffer (50 mM HEPES pH 8, 1% SDS). Samples were sonicated on a probe-less ultrasonic sonicator for five 10-second cycles at 10 watts per tube (Hielscher VialTweeter) to shear genomic DNA. Samples were centrifuged at 18,500 x g to pellet cell debris, and the supernatant was used for subsequent steps. Disulphide bonds were reduced with 5 mM dithiothreitol, followed by 30 min incubation at 60 °C. Free sulfhydryl groups were alkylated by incubating samples in 25 mM iodoacetamide in the dark for 30 min at room temperature. An additional 5 mM of DTT was added to quench the alkylation reaction and samples were incubated at room temperature for 5 minutes. The magnetic bead-based SP3 protocol published in (35) was used to capture proteins prior to digestion. Briefly, magnetic beads were added to proteins in a 10:1 (w/w) ratio. Absolute ethanol was added to bring the ethanol concentration to 70%. Samples were shaken at room temperature for 5 mins at 1000 rpm, and supernatant was discarded. The beads were rinsed 2 times with 80% ethanol, and discarded. Proteins were digested in 100 mM ammonium bicarbonate containing 2 μg of trypsin/Lys-C enzyme mix (Promega) at 37 °C overnight. Peptides were desalted using C18-based solid phase extraction, then lyophilized in a SpeedVac vacuum concentrator. Peptides were solubilized in mass spectrometry-grade water with 0.1% formic acid.

### Shotgun proteomics

Liquid chromatography mass spectrometry/mass spectrometry (LC-MS/MS) data was acquired as previously described (36) with the following modifications: Peptides (2 μg) were loaded on a two-column setup using 2 cm Acclaim PepMap 10 column (75 μm, 3 μm, 100 Å) as trap column and a 50 cm EasySpray ES803 column (75 μm, 2 μm, 100 Å) (Thermo) coupled to an Easy nLC 1000 (Thermo) nano-flow liquid chromatography system connected to Q-Executive mass spectrometer (Thermo). Peptides were separated by reverse phase chromatography using a 265 min non-linear chromatographic gradient of 4-48% buffer B (0.1% FA in ACN) at a flow rate of 250 nl/min. Column temperature was kept at 45 °C. MS data was acquired in positive-ion data-dependent mode. Data-dependent MS analyses were run in a positive top-25 mode. MS1-spectra were acquired for a mass range of *m/z* 350–1800 at a resolution of 140,000, with an automatic gain control (AGC) target of 3□×□10^6^ and 220 ms maximum fill time. The dependent MS/MS spectra were acquired at a resolution of 17,500, with an AGC target of 5□×□10^5^ and 45 ms maximum fill time. The isolation window width was set to 2.0 m*/z*, the isolation offset to 0.4 m*/z* and the intensity threshold to 1.8□×□10^3^. Dynamic exclusion was set to 40 s. Raw data was searched in MaxQuant (37) (version 1.6.3.3) using UniProt protein sequence database containing human protein sequences from Uniprot (complete human proteome; 2019-09). Searches were performed with a maximum of two missed cleavages, and carbamidomethylation of cysteine as a fixed modification. Variable modifications were set as oxidation at methionine and acetylation (N-term). The false discovery rate for the target-decoy search was set to 1% for protein, peptide and site levels. Intensity-based absolute quantification (iBAQ), label-free quantitation (LFQ), and match between runs (matching and alignment time windows set as 0.7 and 20 min respectively) were enabled. The proteinGroups.txt file was used for subsequent analysis. Proteins matching decoy and contaminant sequences were removed, and proteins identified with two or more unique peptides were carried forward. LFQ intensities were used for protein quantitation (38). For proteins with missing LFQ values, median-adjusted iBAQ values were used as replacement (39). Protein intensities were log2-transformed for further analysis. Differentially expressed proteins among the different treatment conditions were identified using Students t-test with multiple test correction using FDR.

### Immunofluorescent microscopy

Cells were seeded onto glass coverslips 24 hours prior to treatments with acute genotoxic stressors. After treatment, cells on coverslips were washed twice with PBS + 0.1% Tween-20 (PBS-T). Cells were fixed for 10 minutes on ice with 3% paraformaldehyde (PFA) + 2% sucrose in PBS. Cells were washed twice in PBS-T, then permeabilized for 10 minutes at room temperature with 0.5% NP-40 solution in PBS. Cells were washed twice in PBS-T, incubated in blocking solution (3% bovine serum albumin [BSA] in PBS-T) for 10 minutes at room temperature. Cells were stained with the appropriate primary antibodies for a given experiment, overnight at 4 °C. The following primary antibodies were used, all at 1:1000 dilution: Phospho-Histone H2AX (Ser139) (Sigma cat #05-636); SF3B1 (Thermo cat #115865). SRSF1 (Thermo cat #324500); cGAS (Cell Signalling Technologies cat #15102S); NUP107 (ThermoFisher cat #PA5-81528); Lamin A/C (Cell Signalling Technologies cat #4777T), Lamin B1 (Abcam cat #ab133741), Emerin (Abcam cat #204987); POLG (Thermo cat #21314); SSBP1 (Proteintech cat #12212); MGME1 (Proteintech cat #23178); CHCHD4 (Bethyl Laboratories cat #A305); CCDC77 (abcam cat #91585); CEP63 (Thermo cat #TA809276S); ATRIP (Thermo cat #TA504642S); UBE2A (Bethyl Laboratories cat #A300). Cells were washed four times in PBS- T, then incubated in fluorescently labelled secondary antibodies for 1 hour at room temperature. The following dilutions were used for secondary antibodies: 1:500 AlexaFluor 568 Goat anti-rabbit IgG secondary (Invitrogen cat #A11036); 1:500 AlexaFluor 488 Goat anti-mouse IgG secondary (Invitrogen cat #A11001); 1:500 AlexaFluor 568 Goat anti-mouse IgG secondary (Invitrogen cat #A11004); 1:500 AlexaFluor 488 Goat anti-rabbit IgG secondary (Invitrogen cat #A11034); Cells were washed four times in PBS-T, then inverted onto a drop of Prolong Glass Antifade Mountant with NucBlue (DAPI) stain (Thermo Fisher Scientific, cat #P36981) on a microscope slide. Slides were imaged on an Olympus immunofluorescent microscope, using CellSens Dimensions software. All IF images presented here were taken at 60X objective on an oil lens. All IF experiments evaluating a percentage of marker-positive MN counted 50 MN per biological replicate. All IF experiments quantifying the abundance of MN counted at least 5 fields of view (FOV) and 200 nuclei per biological replicate. For quantifications of IF intensity presented in Fig. S1C, masks of MN and nuclei were made using the Threshold → Convert to Mask functionality in FIJI (ImageJ) version 2.9.1, and the mean intensity of the area within the masks were calculated using the Analyze → Analyze Particles function. Plot presents the mean pixel intensity of the region within the mask normalized to its area.

### 5-ethynyl uridine (EU) staining

For IF visualization of active transcription in MN, EU staining was carried out using a click-chemistry reaction (Invitrogen cat #C10269) according to manufacturer instructions. In brief: 0.5 mM EU was added to cultured cells, one hour prior to fixation for IF. Cells were fixed and permeabilized as described under the subheading *immunofluorescent microscopy*. Cells were washed twice in PBS-T, then incubated with the click chemistry reaction mixture containing an AlexaFluor 488 reactive azide (Thermo Fisher Scientific, cat #A10266) for 30 minutes at room temperature. Cells were washed twice in PBS-T, then processed as described for immunofluorescent microscopy, beginning from blocking with 3% BSA in PBS-T.

### Data visualization and statistical analysis

For direct comparisons of relative protein abundances between two groups (eg: Fig. 1B-C), two-way *t*-tests using Benjamini-Hochberg (BH)-correction were used to call statistical significance. Clustering analyses among more than two groups of proteins (eg: Fig. 1H, Fig. 3E) was performed by Hartigan-Wong *k*-means clustering, and the number of clusters was chosen by sum of squares. For the assessment presented in Fig. 4, iBAQ measurements for each protein were averaged across the six genotoxic conditions in our panel, from each of the MN and Nuclear samples. We compared protein abundances between MN and nuclei, calling statistically significant differences by Benjamini-Hochberg (BH)-corrected *t*-test (enrichment is statistically greater abundance in MN compared to nuclei, depletion is statistically lower abundance). Next, the original MN protein abundances for each of these significantly enriched or depleted proteins were compared to each of the other enriched or depleted proteins. Only those proteins that had a Pearson correlation coefficient (PCC) of >= 0.7 with at least one other MN protein passed the filter. STRING protein-protein interaction networks and functional enrichment analyses presented in Fig. 2 and Fig. 4 were made in Cytoscape version 3.9.1, where the length of the edges represents the Euclidean distance between nodes from the associated *k*-means clustering analysis (Fig. 2) or the PCC between nodes (Fig. 4). All statistical comparisons and all other visualizations used RStudio version 2022.02.3. Principle component analysis (PCA) used the *prcomp* function from the default *stats* package version 4.2.0 in R. Hierarchical clustering presented in Fig. 2A, Fig, 2E-F, and Fig, 5E used ComplexHeatmap version 2.15.4, with distance method *pearson* and clustering method *complete*. Functional enrichment analysis presented in Fig. 1I, Fig. 3G, Fig. S3E, and Fig. S4D used the *gost* function from the gprofiler2 package version 0.2.1, also available online at https://biit.cs.ut.ee/gprofiler/gost. Queries with the relevant list of differentially expressed proteins (all protein lists used for the functional enrichment analyses in this manuscript are provided in *Supplementary Data)* and the ontology source (KEGG) are indicated on the individual plots. UpSet plots were generated using UpSetR version 1.4.0. All other presented plots were generated using the ggplot function from tidyverse version 1.3.1 in in RStudio version 2022.02.3. All other information regarding biological replicates, sample size, and statistical testing is supplied in the Figure Legends.

## SUPPLEMENTARY INFORMATION TITLES AND LEGENDS

**Supplemental Figure S1. The ionizing radiation (IR)-induced MN proteome recapitulates known MN features. (A)** Volcano plot displaying the log_2_ fold-change (FC) for proteins detected by mass spectrometry (MS), in 10 Gy IR MN compared to nuclei from MCF10A. Log_2_ FC (positive numbers indicating the protein is more abundant in MN) are plotted against the log_10_-transformed Benjamini-Hochberg (BH)-corrected t-test *p*-value (the *q-*value). Differentially expressed proteins are those with a *p*-value < 0.05 and a log_2_ FC greater than 1 or less than -1 (purple colour). Each point represents the mean log_2_ FC for three biological replicates. **(B)** Immunofluorescent (IF) staining for proteins found to be low- or high-abundance in MN compared to primary nuclei by mass spectrometry. 10 Gy IR-exposed MCF10A cells. Representative of three independent experiments. Scale bar = 20 μm. **(C)** Quantification of the fluorescent intensity of each indicated marker in MN compared to primary nuclei by IF staining, as displayed in (B). 10 Gy IR-exposed MCF10A cells. Each point represents a single MN or nucleus. *P-*values indicate result of Student’s t-test. Data from three independent experiments. **(D)** UpSet plot displaying the overlap in detected proteins following MS of 10 Gy IR HeLa-S3 MN compared to MCF10A MN. **(E)** Linear regression correlating the log_2_ fold-change in protein abundance in MN compared to primary nuclei, for 10 Gy IR HeLa compared to MCF10A. Each point represents the mean log_2_ FC value from MCF10A experiments compared to HeLa experiments, each in triplicate. **(F)** The log_2_ fold-change in MN-relevant protein abundance compared to primary nuclei, for 10 Gy IR HeLa compared to MCF10A. **(G)** IF staining for cGAS in unstressed HeLa (top) and MCF10A (bottom). Representative of three independent experiments. Scale bar = 20 μm.

**Supplemental Figure S2. The micronuclear proteome is distinct from the nuclear and cytoplasmic proteomes. (A)** Primary subcellular location for each of the proteins found to be significantly high-abundance in MN samples only (as in Fig. 1H), annotated by the Human Protein Atlas (HPA). Only the top seven populated HPA categories are shown. **(B)** Relative abundance in cGAS protein detected in MN from each of the six stress conditions, by mass spectrometry (MS) in HeLa cells (Z-score). Statistical comparison by one-way ANOVA. **(C-F)** UpSet plots displaying the overlap in the identity of the differentially expressed proteins by stress condition for each of the indicated comparisons (eg. MN compared to cytoplasm). **(G)** Principle component plot displaying the variance among the 54 HeLa samples based on intensity measurements from MS, when the 644 proteins tagged as mitochondrial residents by the HPA are removed from the dataset. **(H)** Relative abundances, measured by MS, of the proteins associated with the GO term “mitochondrial DNA.” Each cell of the heat map represents the mean Z-score from all samples within the indicated subcellular fraction. **(I)** Percentage of MN from MCF10A cells that stained positive for each of the indicated proteins by immunofluorescence (IF). Individual data points presented for IF scoring of MN represent the mean percentage of MN that were positive for the indicated marker, from each biological replicate out of 50 total MN per replicate.

**Supplemental Figure S3. The micronuclear proteome mirrors the proteome of mitotic chromatin. (A-C)** Pearson correlation comparing the relative intensity measurements by mass spectrometry (MS) between our MN dataset and publicly available data describing the proteome of cell cycle-resolved chromatin (25). Z-scores for each protein in the chromatin samples were calculated as described for Fig. 1H. These relative measurements were compared to the MN Z-scores presented in Fig. 1H. **(D)** Consensus clustering by MS intensity measurements. Z-scores were calculated as described for Fig. 1H, using only our MN and the publicly available M-phase chromatin measurements from (C). **(E)** Result of KEGG gene ontology query, where the query list in is the proteins from each of the indicated clusters in (D) and the ∼1,500 proteins shared between our MN samples and the public M-phase chromatin data was used as the background list. Only the top 25 terms (by *p-*value) are shown.

**Supplemental Figure S4. Exogenous genotoxic stress alters both the micronuclear and nuclear protein profiles. (A)** Fingerprint plot displaying the top 100 MN-enriched proteins. Pink squares indicate the protein is more than three times as abundant in MN from the indicated stressor than in MN from any of the other five conditions, making it a putative biomarker candidate. **(B)** Percentage of IR- or DRB-induced MN that stained positive for each of the indicated proteins by immunofluorescence (IF). CCDC77 and CEP63 are putative biomarker candidates for IR and DRB, respectively. ATRIP and UBE2A are proteins that were detected in MN by mass spectrometry (MS), but are not biomarker candidates. **(C)** Volcano plot for the log_2_ fold-change (FC) in proteins detected MS, in pooled MN samples from the five exogenous stress conditions in our panel compared to DMSO MN samples. Log_2_ FC (positive numbers indicating the protein is more abundant in MN) are plotted against the log_10_-transformed Benjamini-Hochberg (BH)-corrected *p*-value (the *q-*value) for the comparison. Differentially expressed proteins are those with a *p*-value < 0.05 and a log_2_ FC greater than 1 or less than -1 (red or black colour). Each point represents the mean log_2_ FC for a single protein across three biological replicates. **(D)** Results of KEGG gene ontology query, where the query list is the proteins significantly depleted in pooled exogenous stress-induced MN compared to DMSO MN (top row) or stress-induced nuclei compared to DMSO nuclei (bottom row). The ∼6,000 total proteins detected in the MN and nuclear samples are the background lists. Only the top 25 terms (by *p-* value) are shown. **(E)** Volcano plot displaying the log_2_ fold-change (FC) in proteins detected by mass spectrometry (MS) in pooled MN samples compared to pooled nuclei. Log_2_ FC (positive numbers indicating the protein is more abundant in MN) are plotted against the log_10_- transformed Benjamini-Hochberg (BH)-corrected t-test *p*-value (the *q-*value). Significantly enriched or depleted proteins have a *q-*value less than 0.05 and a log_2_ FC greater than 1 or less than -1 (green or blue colour). Each point represents the mean log_2_ FC measurement for a single protein across three biological replicates. **(F)** Percent EU+ MN by immunofluorescence (IF) when splicing is perturbed with pladienolide B (PladB) and/or transcription is prevented with actinomycin D (ActD), using the experimental timeline described in Fig. 5D. Statistical comparisons performed using a two-sided Student’s t-test. NS: p=1, ns: *p* > 0.05, *: *p* <= 0.05, **: *p* <= 0.01, ***: *p* <= 0.001, ****: *p* <= 0.0001. All individual points represent the mean percentage of MN that were positive for the indicated marker, from each biological replicate out of 50 total MN per replicate. Error bars represent standard error of the mean, for three independent biological replicates.

