## Supplementary figures and images for "The proteomic landscape of genotoxic stress-induced micronuclei"

### Supplementary Figures

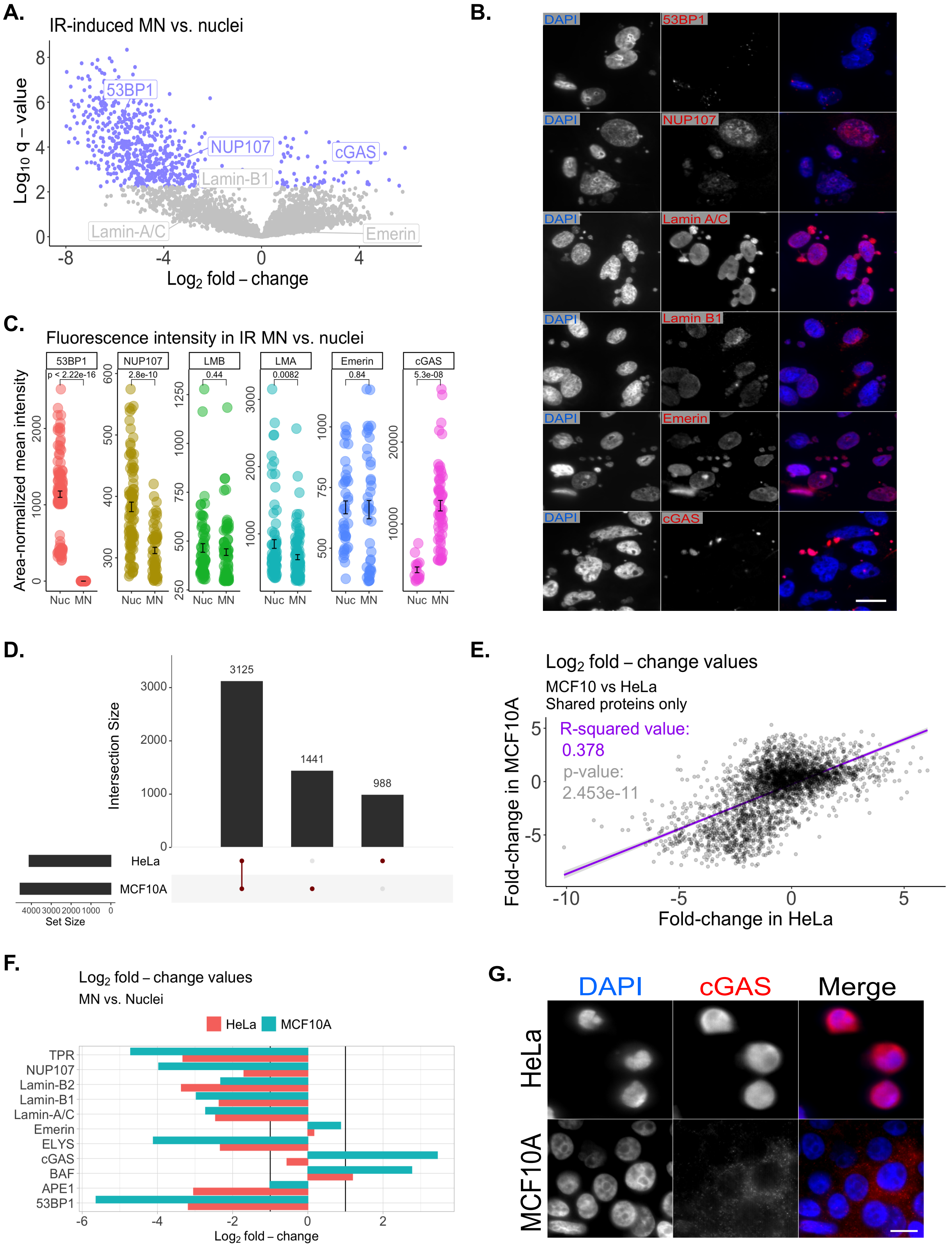

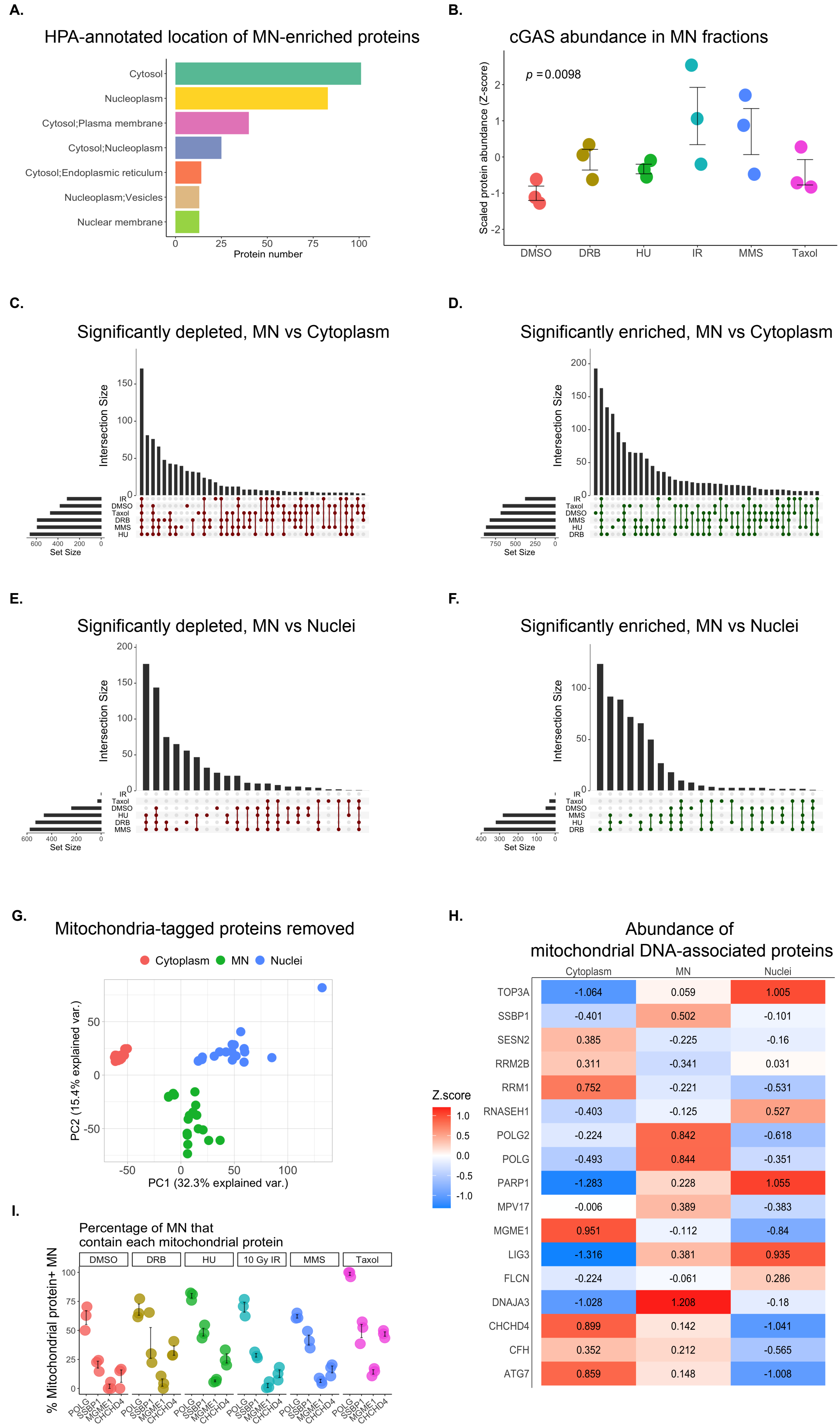

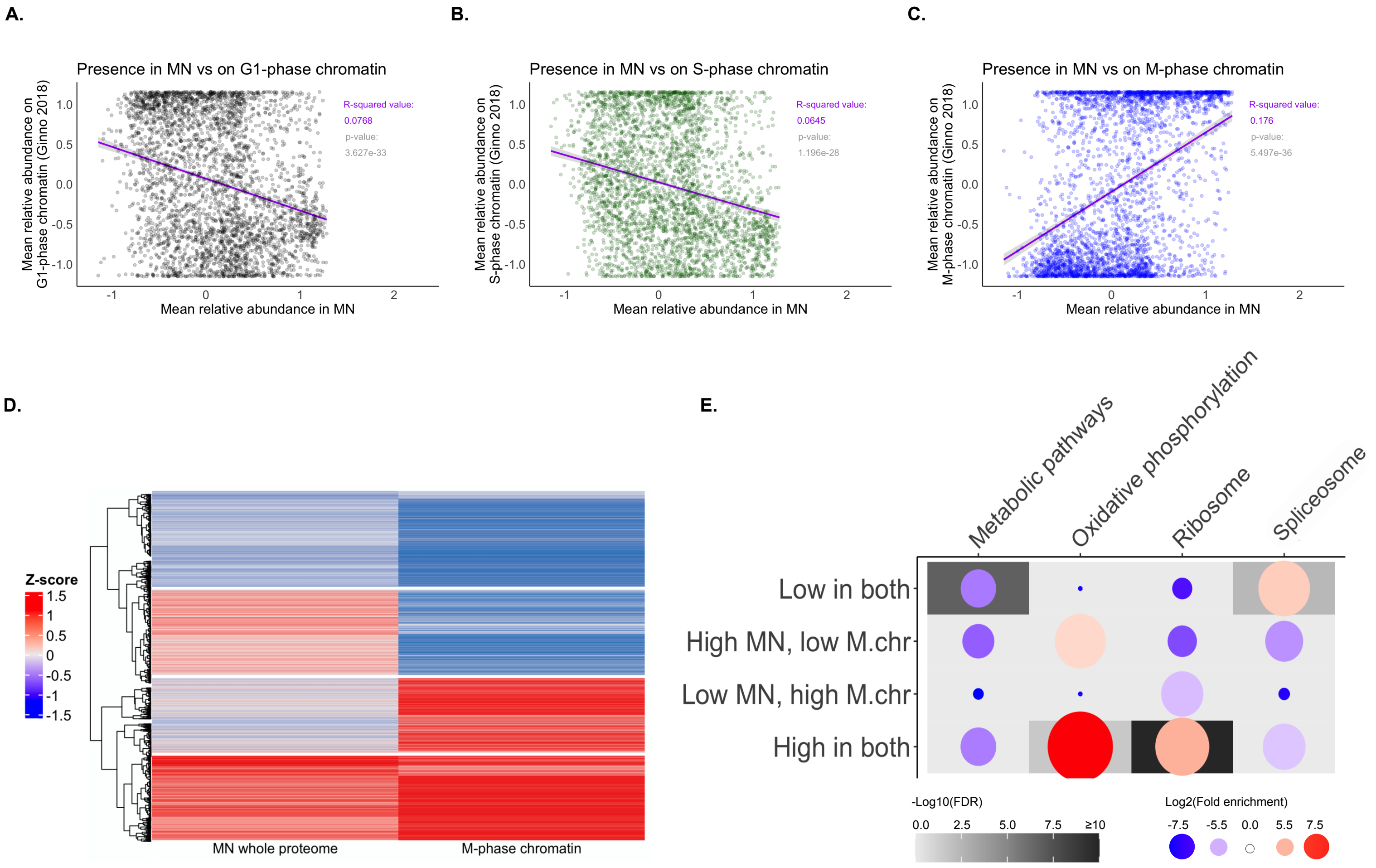

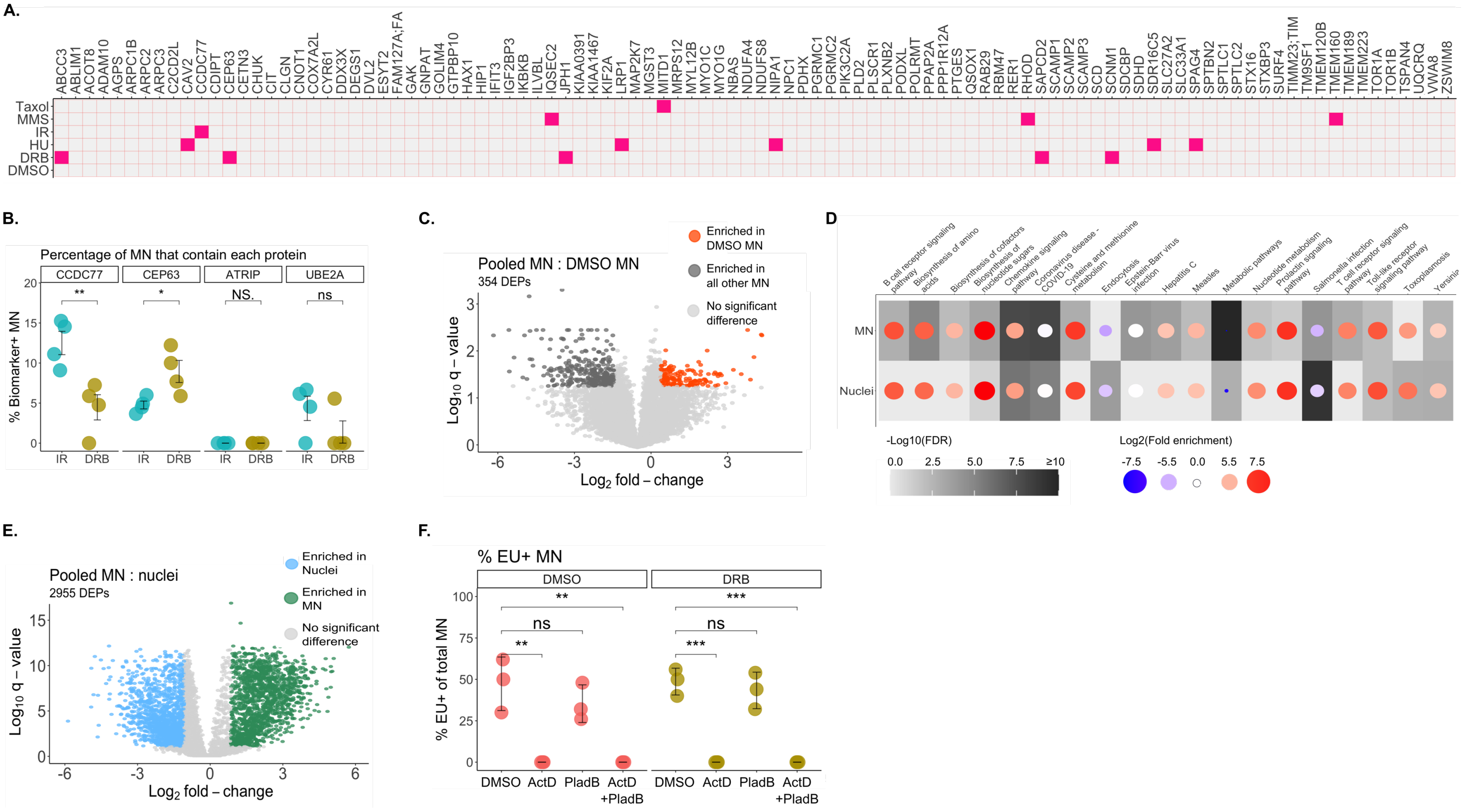
